# Pannexin 1 regulates spiny protrusion dynamics in cortical neurons

**DOI:** 10.1101/2020.03.02.973917

**Authors:** Juan C. Sanchez-Arias, Rebecca C. Candlish, Leigh Anne Swayne

## Abstract

The integration of neurons into networks relies on the formation of dendritic spines. These specialized structures arise from dynamic filopodia-like spiny protrusions. Recently, it was discovered that cortical neurons lacking the channel protein Pannexin 1 (Panx1) exhibited larger and more complicated neuronal networks, as well as, higher dendritic spine densities. Here, we expanded on those findings to investigate whether the increase in dendritic spine density associated with lack of Panx1 was due to differences in the rates of spine dynamics. Using a fluorescent membrane tag (mCherry-CD9-10) to visualize spiny protrusions in developing neurons (at 10 *days-in-vitro*, DIV10) we confirmed that lack of Panx1 leads to higher spiny protrusion density while transient transfection of Panx1 leads to decreased spiny protrusion density. To quantify the impact of Panx1 expression on spiny protrusion formation, elimination, and motility, we used live cell imaging in DIV10 neurons (1 frame every 5 seconds for 10 minutes). We discovered, that at DIV10, lack of Panx1 KO stabilized spiny protrusions. Notably, re-expression of Panx1 in Panx1 knockout neurons resulted in a significant increase in spiny protrusion motility and turnover. In summary, these new data revealed that Panx1 regulates the development of dendritic spines by controlling protrusion dynamics.

**Significance statement:** Cells in the brain form intricate and specialized networks - *neuronal networks* - in charge of processing sensations, executing movement commands, and storing memories. To do this, brain cells extend microscopic protrusions - *spiny protrusions* - which are highly dynamic and survey the local environment to contact other cells. Those contact sites are known as synapses and undergo further stabilization and maturation establishing the function and efficiency of neuronal networks. Our work shows that removal of Panx1 increases the stability and decreases the turnover of spiny protrusion on young neurons.

## Introduction

Pannexin 1 (Panx1) is a four transmembrane domain protein that forms channels permeable to ion and metabolites with various activation mechanisms and diverse (patho)physiological implications (for review Boyce et al., 2018; Chiu et al., 2018). Panx1 is broadly and highly expressed in the brain during post-natal early development (Ray et al., 2005; Vogt et al., 2005) and localized and enriched in synaptic compartments (Sanchez-Arias et al., 2019; Zoidl et al., 2007).

Recent reports have implicated Panx1 in neurite outgrowth, hippocampal synaptic plasticity, and the development of neuronal networks and dendritic spines in cortical neurons (Ardiles et al., 2014; Prochnow et al., 2012; Sanchez-Arias et al., 2019; Wicki-Stordeur & Swayne, 2013). While the behavioural features resulting from a loss of Panx1 have not been thoroughly characterized, a handful of studies have detected important phenotypes like anxiety, increased wakefulness, and spatial learning deficits (Ardiles et al., 2014; Gajardo et al., 2018; Kovalzon et al., 2017; Prochnow et al., 2012). Notably, dendritic spine development has been linked to each of these behaviours. For example, dendritic spine density is increased in various neurodevelopmental disorders in which clinical manifestations include anxiety, intellectual disability, and stereotypical movements (Phillips & Pozzo-Miller, 2015). Moreover, sleep promotes dendritic spine and spiny protrusion turnover in the cortex and hippocampus (Spano et al., 2019; G. Yang & Gan, 2012), which facilitates network sparsity and memory consolidation (Frank et al., 2018; Li et al., 2017). Dendritic spine-based synapses result from spiny protrusions (including dendritic filopodia) actively extending to contact presynaptic boutons during developmental excitatory synaptogenesis; upon contact, spiny protrusions stabilize and evolve into mature dendritic spines along active presynaptic boutons (Fiala et al., 1998; Ziv & Smith, 1996). These steps are critical in establishing network ensembles and Hebbian plasticity (Hoshiba et al., 2017).

## Materials and Methods

In light of this evidence, we investigated the role of Panx1 in spiny protrusion dynamics in cultured primary cortical neurons at 10 *days-in-vitro* (DIV10). We first established an approach to study spiny protrusions using a fluorescent membrane tag (mCherry-CD9-10), allowing us to visualize these characteristically long and thin structures. Then, we transiently transfected wildtype (WT) and Panx1 knock-out (KO) neuronal cultures with EGFP or Panx1EGFP (as well as mCherry-CD9-10) and analyzed spiny protrusions in fixed and living neurons at DIV10. We confirmed that lack of Panx1 leads to higher spiny protrusion density while over-expression and rescue of Panx1 leads to decreased density. Using live cell imaging we observed increased stability and decreased turnover of spiny protrusions in Panx1 KO neurons, while re-expression of Panx1 resulted in a significant increase in spiny protrusion motility and turnover. In summary, these new data reveal an inverse relationship between Panx1 expression and dendritic spine stability.

**Table 1.**
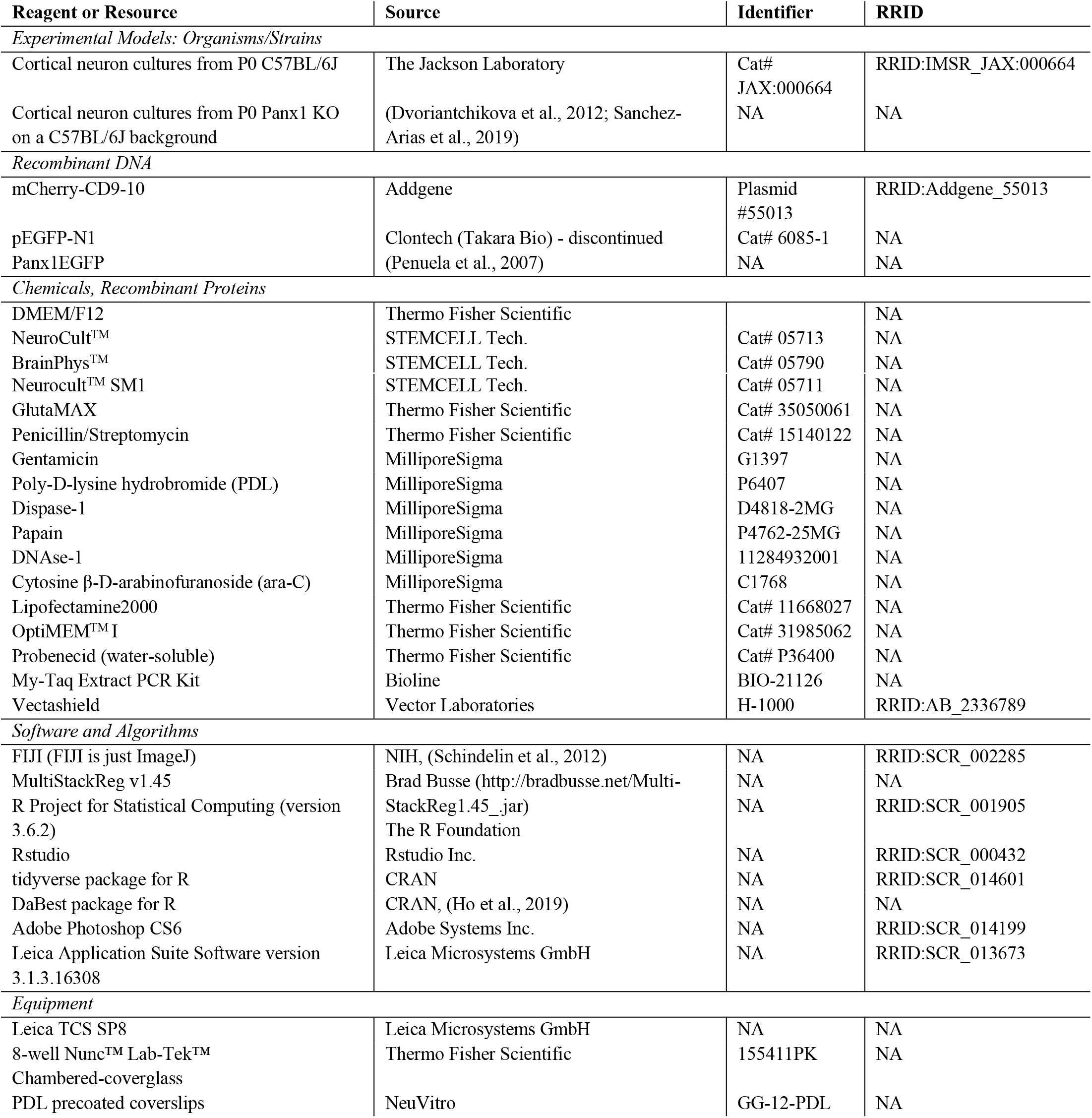
Key Resources Table.

### Experimental animals

All animal procedures were approved by the University of Victoria Animal Care Committee and performed in accordance with the guidelines set by the Canadian Council on Animal Care. Male and female postnatal day (P)0-P1 were used in this study. C57BL/6J mice were obtained from The Jackson Laboratory. The global Panx1 KO strain was derived from a strain originally generated by Dr. Valery Shestopalov (Dvoriantchikova et al., 2012). These mice have been back-crossed in-house onto a C57BL/6J for at least 6 generations (Sanchez-Arias et al., 2019). Mice were housed under a 12 h light/dark cycle starting at 8:00 A.M., with food and water *ad libitum;* temperature was maintained between 20 and 25°C and humidity at 40-65%.

### Primary cortical neuron cultures and transfections

Primary cortical neuron cultures were prepared as previously described (Sanchez-Arias et al., 2019). Briefly, cortices from male and female P0 pups from timed-pregnant WT and Panx1 KO breeding pairs were microdissected and incubated with papain, dis-pase-1, and DNAse-1 for 40 minutes in HBSS followed by mechanical dissociation in DMEM/F12 medium supplemented with Neurocult™ SM1, Gluta-MAX, and penicillin/streptomycin (P/S). Then, 125,000 cells were plated in Nun™ Lab-Tek™ 8-well chambered coverglasses coated with PDL. After 1-2 hours after plating, the medium was replaced with Neurocult™ supplemented with Neurocult™ SM1, GlutaMAX, P/S, and gentamicin. From 4 days-*in-vitro* (DIV) onwards, partial (half) medium changes were done with BrianPhys™ maturation medium (Bardy et al., 2015); to limit proliferation of glial cells, ara-C was added to the medium at DIV4. Transfections were performed at DIV6 using Lipofectamine^©^2000. DNA/lipid complexes were diluted in OptiMEM-I^®^ at ratio of 2 μg DNA:1 μL lipofectamine ratio and incubated at room temperature for 30 minutes. Then, these DNA/lipid complexes were added to cells in BrainPhys™ medium without antibiotics and incubated for 1-1.5 hours. Neurons were transfected with either pEGFP-N1 (250 ng) or Panx1EGFP (250 ng, gift from Silvia Penuela and Dale Laird). All transfections contained mCherry-CD9-10 (250 ng, was a gift from Michael Davidson; Addgene plasmid #55013; http://n2t.net/addgene:55013; RRID:Addgene_55013) to visualize neurons and spiny protrusions (**Figure 1**). All neurons used for this study were used at DIV10. Neurons used for fixed quantifications were plated on PDL-coated coverslips

**Figure 1.**
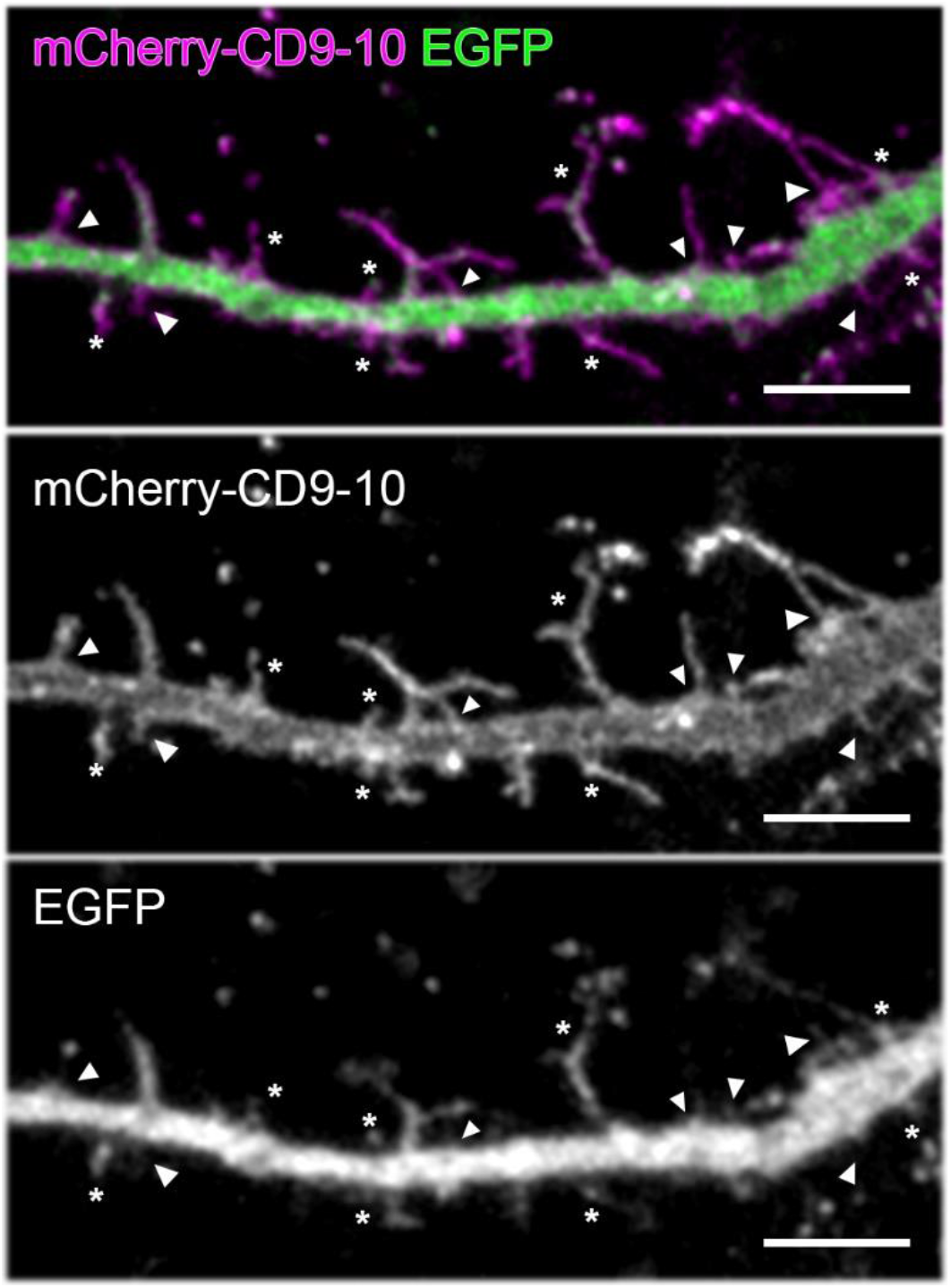
A novel approach to visualize and quantify spiny protrusions in cortical neurons. Representative maximum intensity projection of a dendritic segment from a neuron transfected with mCherry-CD9-10 and EGFP at DIV6 and fixed at DIV10. Thin and long spiny protrusions are more clearly visualized with mCherry-CD9-10 (mid) than the cytoplasmic volume marker EGFP (bottom). Structures not clearly labeled with EGFP are denoted by “*” and those missed entirely are denoted with arrowheads. Scale bar 5 μm.

### Genotyping

Primers for LoxTGF, LoxTGR, and Panx1 LoxR (CTTTGGCATTTTCCCAGTGT, CGCGGTT-GTAGACTTTGTCA, and GTCCCTAC-AGGAGGCACTGA) were used to genotype mice as previously described (Sanchez-Arias et al., 2019). Genomic DNA was extracted from tail-clips using MyTaq™Extract PCR Kit. DNA from WT mice amplifies a single 585 bp band, whereas DNA from global Panx1 KO mice have a single 900 bp band.

### Imaging and analysis of spiny protrusions in fixed cortical neurons

Spiny protrusions (including filopodia) were defined as any membranous protrusions between 0.4 μm and 10 μm. Neurons on coverslips were fixed with 4% PFA and 4% sucrose for 10 minutes and mounted on microscope slides with VectaShield antifade mounting medium. High resolution images (3320×3320, pixel size: 0.088 μm, z-step size: 0.4 μm) were acquired using a Leica TSC SP8 microscope using a 40× immersion oil objective (1.30 NA) and exported to FIJI for analysis (Schindelin et al., 2012). Individual spiny protrusions were traced along the longest neurite (primary neurite) and their density was calculated by dividing the total number of spiny protrusions by the segment length and multiplying by 10 (spiny protrusions per 10 μm). Representative images were processed uniformly with a Gaussian blur of 0.5 pixels, and uniform adjustments to levels and contrast were made using Photoshop CS6 Extended suite (Adobe Systems).

### Imaging and analysis of spiny protrusions in live cortical neurons

Cortical neurons plated on chambered cover-glasses in BrainPhys™ at 37°C and 5% CO2 and pri-mary and secondary dendrite segments of 67-76 μm were imaged (1024×256, pixel size: 0.06 μm) every 5 seconds for 10 minutes and 0.7 μm z-step using a Leica TSC SP8 microscope in resonant mode (8,000 Hz) with a 63× water immersion objective (1.20 NA). Images were exported to FIJI for analysis. First, the four-dimensionality (x,y,z,t) was reduced by creating maximum z projections before additional image processing and x-y drift was corrected with MultiStackReg v1.45 (developed by Brad Busse http://bradbusse.net/MultiStackReg1.45_.jar) when required. Then, images were subjected to a low-pass filter using a Gaussian blur (kernel size 2) and thresholded using the triangle method (Zack et al., 1977). From these binary images, outlines for each time frame were created and temporal colour-coded (**Figure 3*A,B***). Spiny protrusions were manually counted, and four basic characteristics were recorded: formation, elimination, lability, and motility. We defined formation as any *de novo* appearance of a spiny protrusion within the time-lapse recording; elimination was defined as the complete disappearance of a spiny protrusion. Lability was defined as spiny protrusions that were formed and eliminated within the duration of the time-lapse, typically short-lived and lasting 1-3 minutes (**Figure 3*C***). To assess spiny protrusion motility, we annotated partial extensions and partial retractions of individual spiny protrusions (**Figure 3*C***). The survival fraction of spiny protrusions was calculated by dividing the number of spiny protrusions at the end of each time-lapse (10-minute mark) by the number of spiny protrusions at the start (0-minute mark). The overall turnover rate was calculated as the net per cent gain and loss (sum of formation, elimination, and lability) of spiny protrusions divided by the number of spiny protrusions at the start of the time-lapse. Lastly, the overall movement change of spiny protrusions (Δ movement) was calculated by adding the basic dynamic characteristics (formation, elimination, lability, and motility) divided by the number of spiny protrusion at 0 min. Representative images were processed uniformly with a Gaussian blur of 0.5 pixels, and uniform adjustments to levels and contrast were made using Photoshop CS6 Extended suite (Adobe Systems Inc.).

### Experimental design and statistical analysis

For all experiments 3 independent cultures were used. All images were blindly acquired and analyzed. Relevant details are described in *Results*, figure legends, and where appropriate, illustrated on the figures themselves. Data are presented as mean ± standard deviation. Data analysis using bootstrap estimation (5000 bootstrap resamples), determination of effect size, bias-corrected confidence intervals, and Cumming estimation plots were generated using the dabestR package for R (Bernard, 2019; Calin-Jageman & Cumming, 2019; Ho et al., 2019). Null-hypothesis significance testing was performed using R (version 3.6.2) and a *p* value < 0.05 was used as the significant threshold for these tests. Normality was tested using the Shapiro-Wilk test (McDonald, 2014). Group analyses for normally distributed data were performed with a two-way ANOVA coupled to multiple comparisons with Bonferroni’s correction. For non-normally distributed data Kruskal-Wallis pairwise comparisons with Bonferroni’s correction were used.

## Results

**Table 2.**
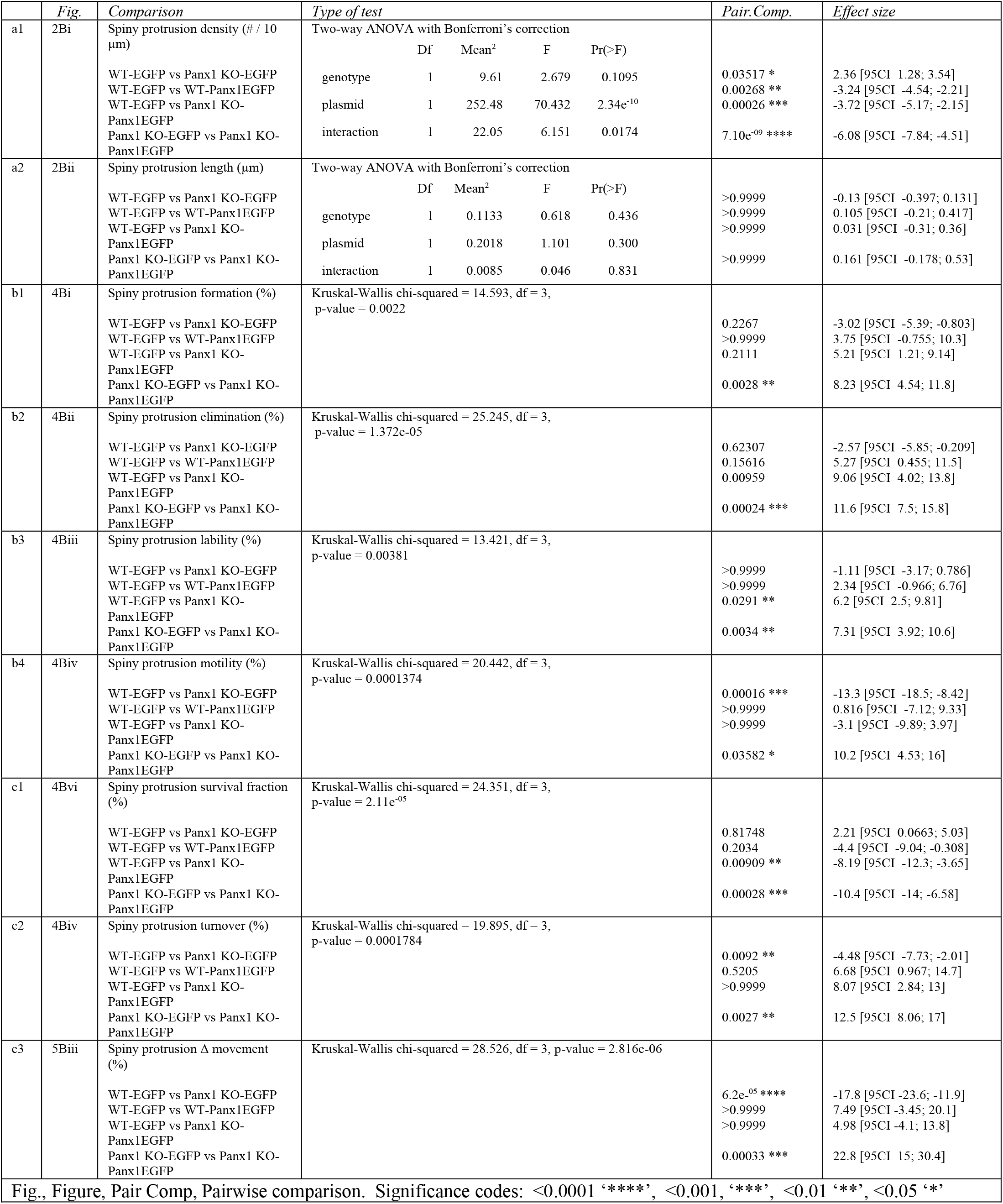
Statistical Table

### A novel approach to visualize and quantify spiny protrusions in cortical neurons

Spiny protrusions (including filopodia) are characteristically highly dynamic, thin, and long. As dendritic arbors mature, these transient structures stabilize into mature dendritic spines. Most methods used to detect these structures rely on cytoplasmic volume markers such as GFP (and its variants) or membranebound lipophilic dyes (DiI, DiO, etc). The former approach allows for sparse labelling but fails to fully label thin processes such a spiny protrusions (**Figure 1**), while the latter achieves clear visualization of these structures by labelling the membrane at the expense of widespread labelling (Mancuso et al., 2013). We transfected cortical neurons with the tetraspanin CD9-10 fused to a monomeric red fluorescent protein mCherry (mCherry-CD9-10 was a gift from Michael Davidson, Addgene plasmid #55013), mCherry-CD9-10 facilitated detailed resolution of spiny protrusions in sparsely transfected cells (**Figure 1**).

### Transfection of Panx1 decreases spiny protrusion density in WT and Panx1 KO DIV10 neurons

To investigate the impact of Panx1 expression, we transfected WT and Panx1 KO cortical neuronal cultures with mCherry-CD9-10 as well as EGFP (control) or Panx1EGFP (over-expression/rescue) at DIV6 and fixed the cells 4 days later at DIV10 (**Figure 2*A***). With EGFP control transfection we observed a 20% increase in spiny protrusion density in primary neurites of Panx1 KO neurons (effect size: 2.36 [95CI 1.28; 3.54], *p* = 0.03517,^a1^). In Panx1EGFP-expressing cultures we observed a 27% decrease in spiny protrusion density in WT neurons (**Figure 2*Bi***, effect size: −3.24 [95CI −4.54; −2.21],*p* = 0.00268,^a1^) and a 42.5% density reduction in Panx1 KO neurons (**Figure 2*Bi***, effect size: −6.08 [95CI – 7.84; −4.51], *p* <0.0001,^a1^). Spiny protrusion length was not significantly different amongst the groups (**Figure 2B*ii***,^a2^). These results suggest spiny protrusion density is inversely proportional to Panx1 expression levels.

**Figure 2.**
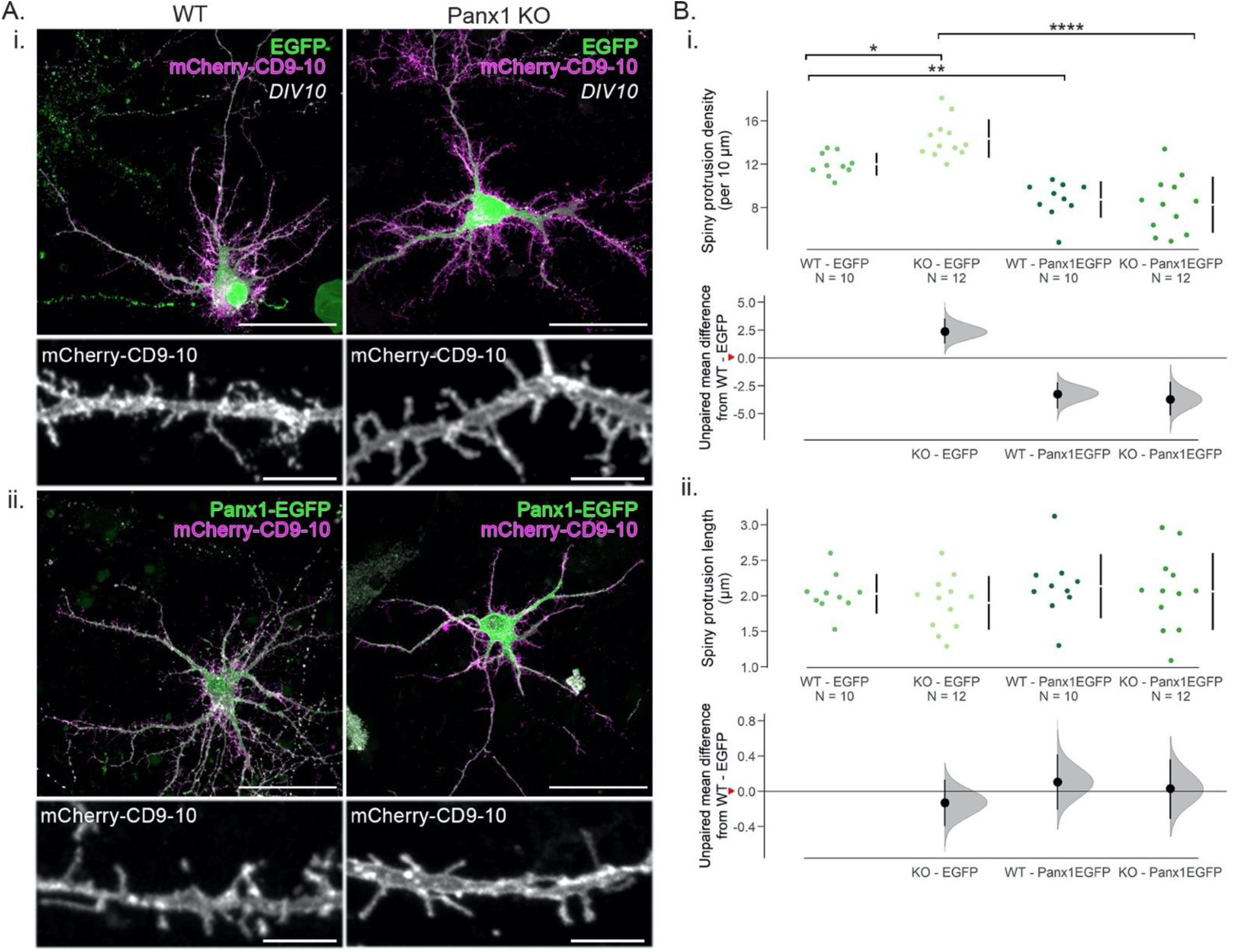
Spiny protrusion density is inversely related to Panx1 expression levels in DIV10 neurons. ***A**.* Representative maximum intensity projections of WT and Panx1 KO cultured cortical neurons transfected with mCherry-CD9-10 and either EGFP *(**Ai**)* or Panx1EGFP (***Aii***) as well as cropped images of their respective dendritic segments from a primary neurite. Scale bar: 50, and 5 μm. ***B***. Effect of Panx1 expression in spiny protrusion density and length in developing cortical neurons transfected with mCherry-CD-9-10 and either EGFP or Panx1EGFP using Cumming estimation plots. ***Bi***. With EGFP expression, spiny protrusion density was higher with Panx1 KO neurons (WT-EGFP: 12.0 ± 0.3 spiny protrusions per 10 μm; Panx1 KO-EGFP: 14.4 ± 0.5 spiny protrusions per 10 μm, *p* = 0.03517, two-way ANOVA with Bonferroni’s multiple-comparison test). With Panx1EGFP expression, spiny protrusion density was decreased in both WT and Panx1 KO neurons (WT-Panx1EGFP: 8.8 ± 0.5 spiny protrusions per 10 μm, *p* = 0.00268; Panx1 KO-Panx1EGFP: 8.3 ± 0.8 spiny protrusions per 10 μm, *p* < 0.0001, two-way ANOVA with Bonferroni’s multiple-comparison test, ^a1^). *Bii.* No significant differences in spiny protrusion length were found between groups (WT-EGFP: 2.0 ± 0.3 μm; Panx1 KO-EGFP: 1.9 ± 0.4 μm, *p* > 0. 9999, two-way ANOVA with Bonferroni’s multiple-comparison test, ^a2^; WT-Panx1EGFP: 2.1 ± 0.1 μm; Panx1 KO-Panx1EGFP: 2.1 ± 0.2 μm, *p* > 0. 9999, two-way ANOVA with Bonferroni’s multiple-comparison test). Data are presented as mean ± standard deviation. Effect sizes are reported in the main text and Table 2. Red arrowheads on the y-axis on the bottom panel of Cumming estimation plots represent WT-EGFP means. s.p., spiny protrusion. <0.0001, ‘****’; <0.001, ‘***’; <0.01 ‘**’; <0.05 ‘*’. *Figure on next page.*

### Measuring spiny protrusion dynamics in living neurons using a membrane marker

To investigate the mechanisms contributing to differences in spiny protrusion densities between groups, we acquired 10-minute time-lapses (one frame every 5 seconds) of primary and secondary dendrites from cortical neurons at DIV10. These cultures were transfected with mCherry-CD9-10 and either EGFP or Panx1EGFP at DIV6. At DIV10, dendrites harbour highly dynamic, thin, and long spiny protrusion that are the precursors for dendritic spines (Fiala et al., 1998; Ziv & Smith, 1996). We reduced the dimensionality of the time-lapses by creating maximum z-projections, and then images were passed through a low-pass filter and thresholded to create outlines (**Figure 3*A***). The dendritic silhouettes (**Figure 3*B***) were then temporally colour-coded to facilitate the detection of formation, elimination, liability, retraction, and growth of spiny protrusions (**Figure 3*C***).

**Figure 3.**
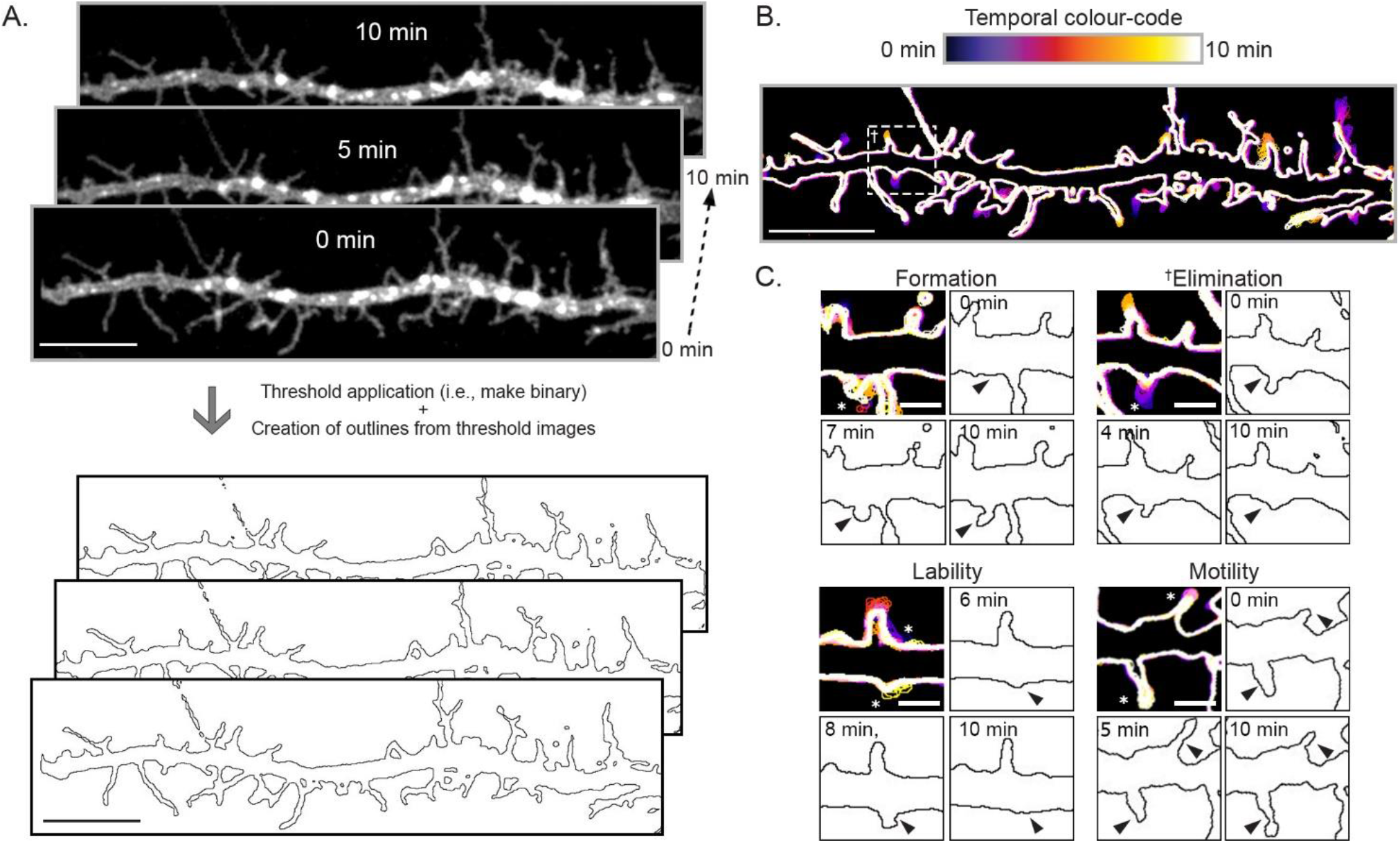
Image analysis strategy to quantify spiny protrusion dynamics in cortical cultures. Ten minutes time-lapses were acquired by imaging dendrite segments from cortical neurons every 5 seconds. Note that this a DIV10 WT cortical neuron transfected with mCherry-CD9-10 and Panx1EGFP; only mCherry-CD9-10 is shown. The dimensionality of these recordings was reduced by creating maximum z projections. Images were thresholded to create outlines (***A***, scale bar 10 μm), which were temporally colour-coded (***B***, scale bar 10 μm), allowing the visualization of various events such as the percentage of spiny protrusion (relative to time 0) undergoing formation (*de novo* appearance), elimination (complete disappearance by the end of the time-lapse), lability (appearance and disappearance by the end of the time-lapse), and retraction/extension (incomplete shrinkage or growth to an existing protrusion) shown in (***C***, scale bar 2 μm). Note that examples in ***C*** (cropped to show highlight the event in question) come from different cultures and different genotypes all at DIV10 transfected with mCherry-CD9-10 and either EGFP or Panx1EGFP at DIV6. The example for elimination (**†**) comes from the neurite in ***B***. Data in **Figure 4** includes quantification of these examples. See Methods for further details.

### Basic characteristics of spiny protrusion dynamics in WT and Panx1 KO neurons at DIV10

Using the above approach, we observed that transfection of Panx1EGFP in Panx1 KO neurons significantly increased the percentage of formation and elimination of spiny protrusions compared to EGFP transfection of Panx1 KO neurons (**Figure 4*A* & 4*Bi-ii***, formation – effect size KO-EGFP *vs.* KO-Panx1EGFP: 8.23% [95CI 4.54%; 11.8%], *p* = 0.0028, ^b1^; elimination – effect size KO-EGFP *vs.* KO-Panx1EGFP: 11.6% [95CI 7.5%; 15.8%], *p* = 0.00024 ^b2^), while no significant differences were observed between Panx1EGFP and EGFP transfection in WT neurons (*p* > 0.9999, ^b1^). Similarly, no significant differences were observed between genotypes with EGFP (control) transfection (**Figure 4*Bi-ii***, formation – effect size WT-EGFP *vs.* KO-EGFP: −3.02% [95CI −5.39%; −0.803%], *p* = 0.2267, ^b1^; elimination – effect size WT-EGFP *vs.* KO-EGFP: −2.57% [95CI −5.85%; −0.209%], *p* = 0.62307, ^b2^). We next quantified spiny protrusion lability within our experimental groups. Transient expression of Panx1EGFP in Panx1 KO neurons significantly increased spiny protrusion lability; there were no significant effects of Panx1EGFP expression in WT neurons (**Figure 4*Biii***, effect size KO-EGFP *vs.* KO-Panx1EGFP: 7.31% [95CI 3.92%; 10.6%], *p* = 0.0034; effect size WT-EGFP *vs.* WT-Panx1EGFP: 2.34% [95CI −0.966%; 6.76%], *p* > 0.9999, ^b3^). There was also no significant effect of EGFP expression between WT and Panx1 KO neurons (effect size WT-EGFP *vs.* KO-EGFP: −1.11%[95CI −3.17%; 0.786%], *p* >0.9999, ^b3^). Additionally, within groups transiently expressing EGFP, Panx1 KO neurons exhibited significantly reduced spiny protrusion motility (**Figure 4*Biv***, effect size WT-EGFP *vs.* KO-EGFP: −13.3% [95CI −18.5%; −8.42%], *p* = 0.00016, ^b4^). Intriguingly, transient Panx1EGFP expression increased spiny protrusion motility in Panx1 KO neurons only (effect size KO-EGFP *vs.* KO-Panx1EGFP: 10.2% [95CI 4.53%; 16%], *p* = 0.03582, ^b4^). Together these results suggest that spiny protrusion dynamics roughly correlate with Panx1 expression levels.

**Figure 4.**
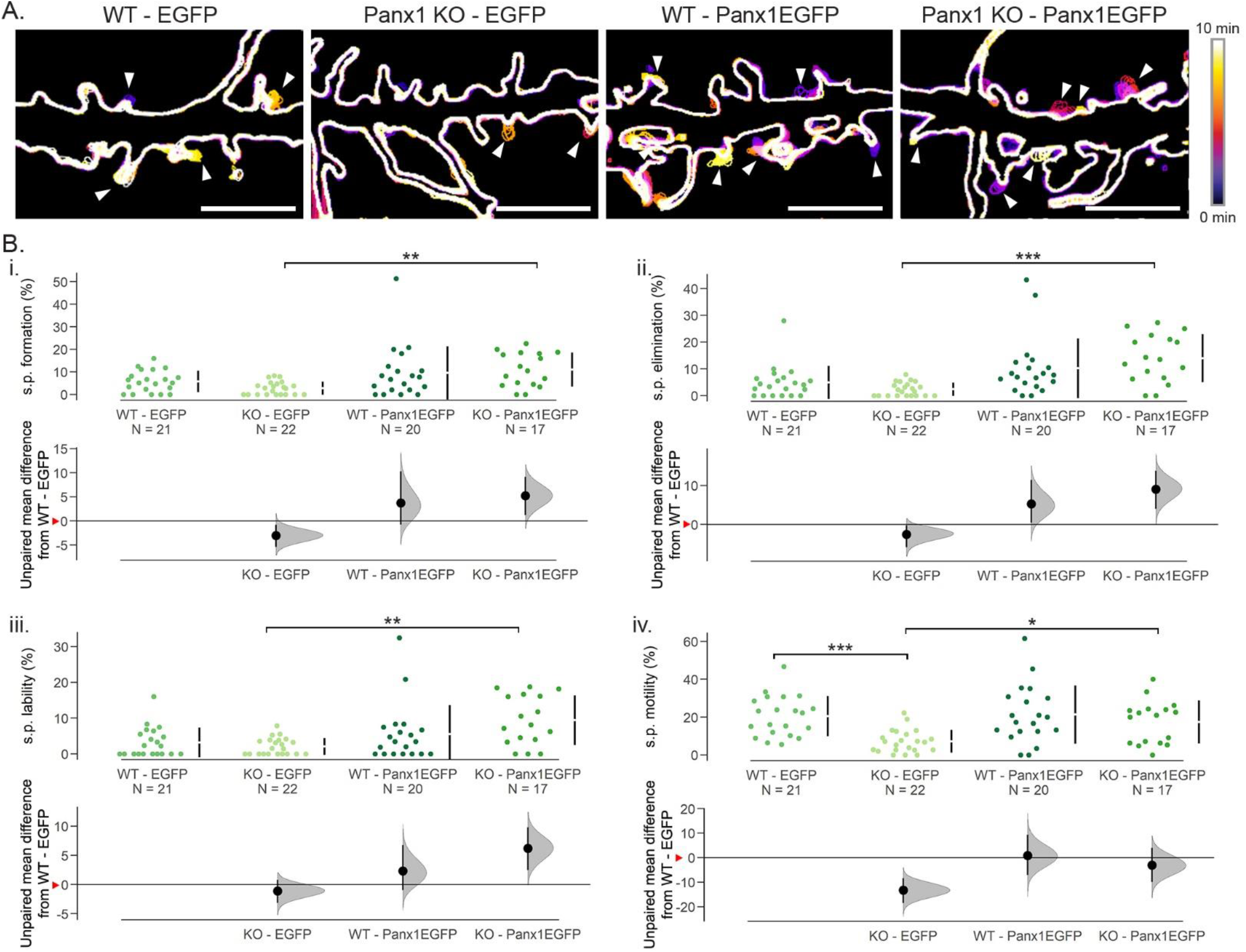
Basic characteristics of spiny protrusion dynamics in WT and Panx1 KO neurons at DIV10. ***A**.* Representative colour-coded outlines of WT and Panx1 KO neurons transfected with mCherry-CD9-10 and either EGFP or Panx1EGFP showing examples of spiny protrusion formation, elimination, lability, and motility events (arrowheads). These examples are cropped from the full regions of analysis from primary neurites. ***B***. Effect of Panx1 expression on spiny protrusion formation, elimination, lability, and motility in WT and Panx1 KO using Cumming estimation plots. ***Bi***. Spiny protrusion formation was significantly higher in Panx1 KO neurons transiently expressing Panx1EGFP compared to those expressing EGFP (KO-EGFP: 0.2% ± 0.1%, KO-Panx1EGFP: 4.6% ± 1.3%, *p* = 0.0028, Kruskal-Wallis test, ^b1^). No significant differences were observed between genotypes in EGFP-expressing neurons (WT-EGFP: 1.7% ± 0.7%; Panx1 KO-EGFP: 0.2% ± 0.1%, *p* = 0.2267, Kruskal-Wallis test, ^b1^). ***Bii**.* Similarly, only transient expression of Panx1EGFP in Panx1 KO neurons increased spiny protrusion elimination (KO-EGFP: 0.3% ± 0.15%; KO-Panx1EGFP: 4.6% ± 1.28%, *p* = 0.00024, Kruskal-Wallis test, ^b2^). No significant differences were found between WT and Panx1 EGFP-expressing cells *(p* = 0.62307, ^b2^). ***Biii*** Spiny protrusion lability was higher in Panx1 KO neurons transfected with Panx1EGFP (KO-EGFP: 2.1% ± 0.5%; KO-Panx1EGFP: 9.4% ± 1.7%, *p* = 0.0034, Kruskal-Wallis test, ^b3^), beyond that observed in WT expressing EGFP control *(p* = 0.0291, Kruskal-Wallis test, ^b3^). Transient expression of Panx1EGFP in WT neurons had no significant effects *(p* >0.9999, ^b3^). ***Biv**.* Spiny protrusion motility was significantly reduced in Panx1 KO neuron expressing EGFP control (WT-EGFP: 20.5% ± 2.3%; KO-EGFP: 7.2% ± 1.3%, *p* = 0.00016, Kruskal-Wallis test, ^b4^). Transient Panx1EGFP expression increased spiny protrusion motility in Panx1 KO neurons only (KO-Panx1EGFP: 17.4% ± 2.8%, *p* = 0.03582, Kruskal-Wallis test, ^b4^). Effect sizes are reported in the main text and Table 2. Red arrowheads on the y-axis on the bottom panel of Cumming estimation plots represent WT-EGFP means. s.p., spiny protrusion; <0.001, ‘***’; <0.01 ‘**’; <0.05 ‘*’.

### Panx1 KO neuron spiny protrusions are more stable

We used the basic characteristic measurements devised in Figure 3C to calculate spiny protrusion survival fraction, turnover, and overall change in movement (Δ movement). The number of spiny protrusions persisting at the end of the analysis period relative to time 0 min, referred to as survival, was significantly reduced in Panx1EGFP expressing Panx1 KO neurons (**Figure 5*A* & 5*Bi***, KO-EGFP *vs.* KO-Panx1EGFP: −10.4% [95CI −14%; −6.58%], *p* = 0.00028, ^c1^). This reduction of survival surpassed that seen in WT, EGFP expressing cells (effect size WT-EGFP *vs.* KO-Panx1EGFP: −8.19% [95CI −12.3%; −3.65%], *p* = 0.00909, ^c1^). We next calculated turnover by adding together formation, elimination, and lability, divided by the total number of spiny protrusions at time 0 min. With EGFP expression, turnover was significantly reduced in Panx1 KO neurons (**Figure 5*Bii***, effect size WT-EGFP *vs.* KO-EGFP: −4.48% [95CI −7.73%; −2.01%], *p* = 0.0092, ^c2^). Within Panx1 KO cultures transient Panx1EGFP expression significantly increased turnover compared to EGFP control (effect size KO-EGFP *vs.* KO-Panx1EGFP: 5.24% [95CI 2.87%; 8.66%], *p* = 0.0027, ^c2^). Finally, to measure the overall change in spiny protrusion movement (Δ movement), we calculated the sum of the four basic dynamic characteristics (formation, elimination, lability, and motility). Within EGFP expressing cells, Δ movement was significantly reduced in Panx1 KO neurons compared to WT controls (**Figure 5*Biii panel***, effect size WT-EGFP *vs.* KO-EGFP: −17.8 [95CI −23.6; −11.9], *p* < 0.0001). Transient Panx1EGFP expression resulted in increased Δ movement in Panx1 KO cultures only (effect size KO-EGFP *vs.* KO-Panx1EGFP: 22.8 [95CI 15; 30.4], *p* = 0.00033, ^c3^). Altogether, these results suggest that Panx1 KO neuron spiny protrusions are more stable.

**Figure 5.**
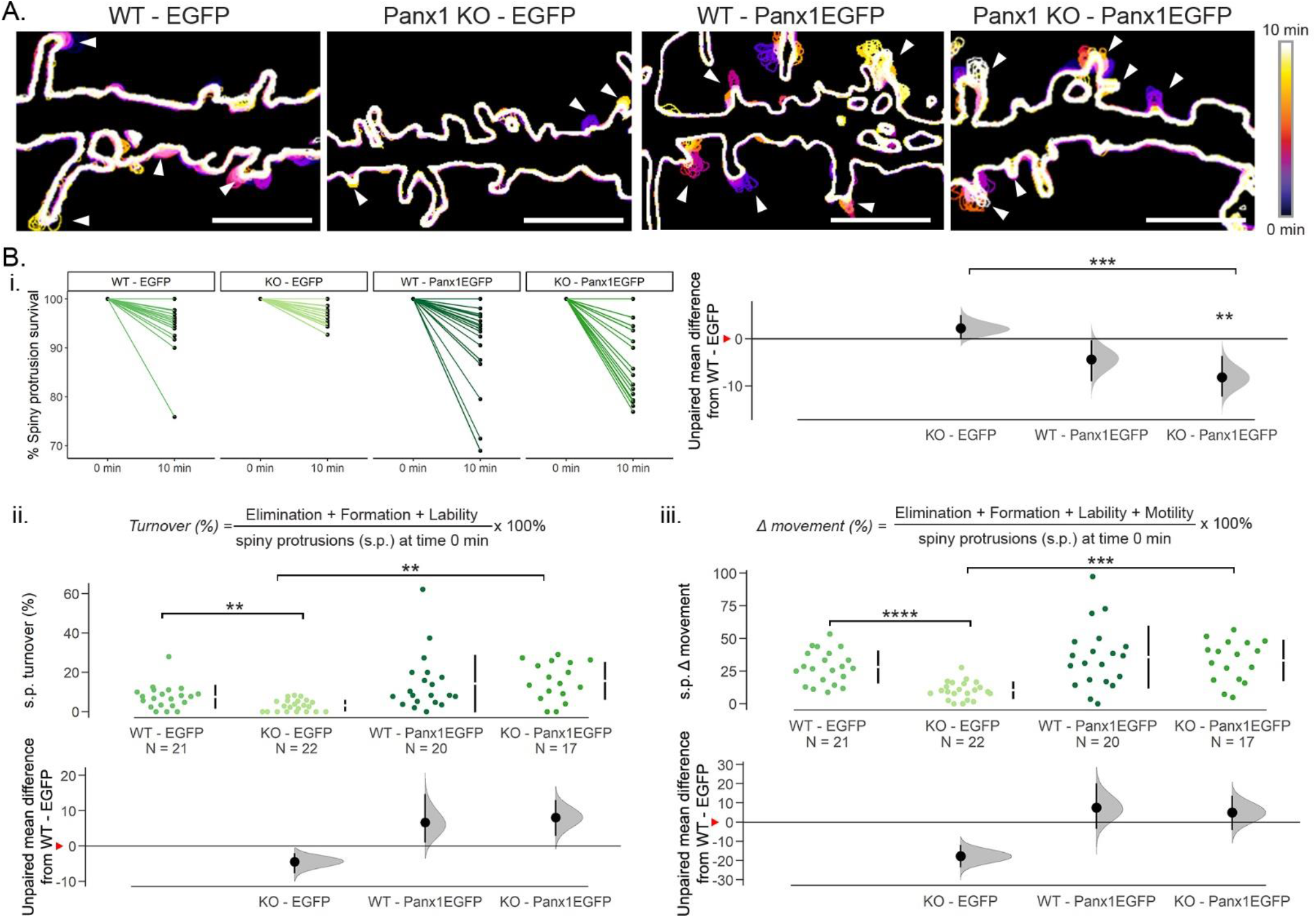
Panx1 KO neuron spiny protrusions are more stable. ***A.*** Representative colour-coded outlines of WT and Panx1 KO neurons transfected with mCherry-CD9-10 and either EGFP or Panx1EGFP showing examples of spiny protrusion movement (arrowheads). These examples are cropped from the full regions of analysis from primary neurites. ***B.*** Cumming estimation plots of spiny protrusion second order metrics: survival fraction, turnover, and overall change in movement (Δ movement). ***Bi.*** Transient Panx1 expression in WT and Panx1 KO neurons decreased the survival fraction of spiny protrusions; however, this was only statistically significant in Panx1 KO neurons (WT-EGFP: 94.5 ± 1.2%; WT-Panx1EGFP: 91.1% ± 1.9%, *p* = 0.2034, ^c1^; Panx1-EGFP: 97.7% ± 0.5%; Panx1 KO-Panx1EGFP: 87.3% ± 1.9%, *p* = 0.00028, Kruskal-Wallis test, ^c1^). ***Bii.*** In the EGFP-control-expressing group, spiny protrusion turnover was reduced in Panx1 KO neurons (WT-EGFP: 7.5% ± 1.3; Panx1-EGFP: 3.1% ± 0.6%, *p* = 0.0092, Kruskal-Wallis test, ^c2^). Transient expression of Panx1 significantly increased spiny protrusion turnover in Panx1 KO neurons but not in WT neurons (WT-Panx1EGFP: 14.2% ± 3.3%,*p* > 0.9999; Panx1 KO-Panx1EGFP: 15.6% ± 2.34%,*p* = 0.0027, Kruskal-Wallis test, ^c2^). ***Biii.*** Spiny protrusion overall movement change (Δ movement) was reduced in Panx1 KO neurons (WT-EGFP: 28% ± 2.8%; KO-EGFP: 10.3% ± 1.5%, *p* <0.0001, Kruskal-Wallis test, ^c3^). Panx1EGFP expression increased Δ movement in both WT (WT-Panx1EGFP: 35.5% ± 5.4%) and Panx1 KO neurons; however, this effect was only significant in Panx1 KO neurons (KO-Panx1EGFP: 33% ± 3.8%, *p* = 0.00033, Kruskal-Wallis test, ^c3^). Effect sizes are reported in the main text and Table 2. Red arrowheads on the y-axis on the bottom panel of Cumming estimation plots represent WT-EGFP means. s.p., spiny protrusion. <0.0001, ‘****’; <0.001, ‘***’; <0.01 ‘**’.

## Discussion

Dendritic spine-based synapses account for the bulk of excitatory neurotransmission in the cerebral cortex and have been implicated in neurodevelopmental and neuropsychiatric disorders (Forrest et al., 2018; Kwon et al., 2019; Lima-Caldeira et al., 2019; Nishiyama, 2019). Although the mechanisms underlying plasticity of existing dendritic spines have been well characterized (Araya et al., 2014; Holtmaat et al., 2005; Sala & Segal, 2014; Schätzle et al., 2018), the processes involved in their formation are less well understood (Sando et al., 2017; Sigler et al., 2017; reviewed in Südhof, 2018). Here we identified a novel role for Panx1 in regulating the dynamics of developing dendritic spines, building on previous work showing that Panx1 KO cortical neurons exhibit higher dendritic spine density and more complex networks. Our results found a reciprocal relationship between Panx1 expression levels and spiny protrusion stability. Additionally, in order to make these discoveries, we optimized methods for the visualization and analysis of spiny protrusion dynamics, thereby providing a framework for others. While the current study is limited to a single timepoint (DIV10), our in-triguing results suggest that additional longitudinal analysis is now warranted. Although *in vitro* spine plasticity characteristics correlate highly with those observed in more complex models (e.g. slice), it would be useful to confirm the current observations within these systems.

The relatively muted impact of Panx1EGFP over-expression in WT neurons, suggests that Panx1 effect on spiny protrusion dynamics is subject to saturation. Possible mechanisms of saturation could be limited machinery for trafficking supplementary Panx1 to spiny protrusions or self-regulation via ATP-dependent internalization. Alternatively, the effects of supplementary Panx1 could be constrained by limited amounts of endogenous interacting partners, such as Crmp2, Arp3c, and actin or saturation of downstream autocrine or paracrine purinergic signalling pathways related to its ATP release function (e.g. ATP stimulating glia) (Abbracchio et al., 2009; Bhalla-Gehi et al., 2010; Dahl, 2015; Wicki-Stordeur & Swayne, 2013, p.; Xu et al., 2018; D. Yang et al., 2015). In contrast to WT neurons, transient expression of Panx1EGFP exhibited significant effects on spiny protrusion dynamics in Panx1 KO cultures. Somewhat consistent with previous results, in EGFP-control-expressing cultures, loss of Panx1 precociously stabilized spiny protrusions, pointing to a fundamentally different underlying molecular organization.

Consistent with this idea, recent work has identified brain enriched and autism associated single nucleotide polymorphisms (SNPs) resulting in changes in Panx1 expression levels; although the direction of this change (i.e. decrease or increase Panx1 expression) was not identified (Davis et al., 2012). Further supporting a role for Panx1 in neuronal development, intellectual disability was observed in an individual with a germline single nucleotide polymorphism in PANX1 (Shao et al., 2016).

In addition to playing a direct role in neurodevelopment, Panx1 is also indirectly involved through its interaction with Crmp2 and purinergic receptor signalling (Boyce et al., 2015; Boyce & Swayne, 2017; reviewed in Swayne & Boyce, 2017). Crmp2 auto-antibodies have been implicated in ASD (Braunschweig et al., 2013), while suramin treatment corrected synaptic and behavioural phenotypes in the Fragile X mouse model (J. C. Naviaux et al., 2015; R. K. Naviaux et al., 2013, 2017).

In summary, this work significantly advances our understanding of the role on Panx1 in dendritic spine development and underscores the importance of additional molecular mechanistic studies investigating intrinsic (e.g. Crmp2) and extrinsic (e.g. glia) pathways.

## Acknowledgements

We are thankful for technical assistance from Sarah N. Ebert who was supported by a Jamie Cassels Undergraduate Research Award. We are also thankful for assistance from Reg Sidhu (Leica Microsystems) in the optimization of our live imaging set up.

## Notes

**Funding sources**: This work was supported by operating grants from the Canadian Institutes of Health Research (CIHR Grant MOP142215), from the Natural Sciences and Engineering Research Council (NSERC) [RGPIN-2017-03889], The Scottish Rite Charitable Foundation of Canada (15118) and the University of Victoria-Division of Medical Sciences to L.A.S. L.A.S. was also supported by a Michael Smith Foundation for Health Research and British Columbia Schizophrenia Society Foundation Scholar Award (5900). J.C.S.A. was supported by a University of Victoria Fellowship Graduate Award. L.A.S. is also grateful for infrastructure support from the Canada Foundation for Innovation (29462) and the BC Knowledge Development Fund (804754) for the Leica SP8 confocal microscope system.

#### Summary of Updates

We updated the acknowledgements section and updated Figures 3, 4, & 5 as well as all the figure legends.

